# Bidirectional introgression between *Betula tianshanica* and *Betula microphylla* and its implications for conservation

**DOI:** 10.1101/2020.06.05.135285

**Authors:** Junyi Ding, Donglai Hua, James S. Borrell, Richard J.A. Buggs, Luwei Wang, Feifei Wang, Zheng Li, Nian Wang

## Abstract

- Molecular markers can allow us to differentiate species that occupy a morphological continuum, and detect patterns of allele sharing that can help us understand the dynamics of geographic zones where they meet. *Betula microphylla* is a declining wetland species in NW China that forms a continuum of leaf morphology with its relative *Betula tianshanica*.
- We use ecological niche models (ENM) to predict the distribution of *B. microphylla, B. tianshanica* and the more commonly occurring *B. platyphylla*. We use restriction-site associated DNA sequencing and SSRs to resolve their genetic structure and patterns of allele sharing.
- ENM predicted an expansion of suitable range of *B. tianshanica* into *B. microphylla* since the Last Glacial Maximum and the contraction of *B. microphylla’s* range in the future. We resolved the species identification of some intermediate morphotypes. We found signatures of bidirectional introgression between *B. microphylla* and *B. tianshanica* with SNPs showing more admixture than SSRs. Introgression from *B. microphylla* into *B. tianshanica* was greater in the Tianshan Mountains where the two species have occurred in proximity. Unexpectedly, introgression from *B. tianshanica* into *B. microphylla* was widespread in the Altay Mountains where there are no records of *B. tianshanica* occurrence.
- This presence of *B. tianshanica-derived* alleles far beyond the species’ current range could be due to unexpectedly high pollen flow, undiscovered populations of *B. tianshanica* in the region, incomplete lineage sorting, or selection for adaptive introgression in *B. microphylla*. These different interpretations have contrasting implications for the conservation of *B. microphylla*.

## 1. INTRODUCTION

Hybridisation can play an important role in plant evolution by introducing novel alleles (Goulet et al. 2017) and leading to speciation (Abbott et al. 2013; Whitney et al. 2010). However, hybridisation can also lead to significant conservation challenges where it involves rare and common species (Balao et al. 2015), native and invasive species (Bleeker et al. 2007), or a domesticate and its wild ancestor (Wayne and Shaffer 2016). Hybridisation can potentially reduce fitness via genetic or demographic swamping, which may introgress maladaptive alleles (Rhymer and Simberloff 1996; Todesco et al. 2016). Such a situation may be worse for wind-pollinated tree species because pollen may transfer long distances (Hjelmroos 1991; Skjøth et al. 2007), leading to hybridisation far beyond the current species’ distribution.

The genus *Betula* is an excellent model to assess the consequences of hybridisation among species. As a genus consisting of approximately 65 species and subspecies (Ashburner and McAllister 2016; Wang et al. 2016), extensive hybridisation and introgression among *Betula* species have been documented based on morphology, cytogenetics and molecular markers (Anamthawat-Jónsson and Thórsson 2003; Anamthawat-Jónsson and Tómasson 1990; Bona et al. 2018; Eidesen et al. 2015; Hu et al. 2019; Tsuda et al. 2017). *Betula* species are pioneers, colonising new habitats and providing shelter for other trees. The fact that species of *Betula* often co-occur and are wind-pollinated means that hybridisation is frequent among them (Ashburner and McAllister, 2016). Some species have restricted distributions, such as *B. nana* in the UK, possibly due to historical hybridisation and habitat fragmentation (Borrell et al. 2018; Wang et al. 2014; Zohren et al. 2016). Several others are rare or endangered, such as *B. megrelica* from Georgia and *B. calcicola* from southwest China (Ashburner and McAllister, 2016).

In this study, we investigate the possible role of hybridisation and introgression in the decline of the regional endemic tree *B. microphylla*, which is on the verge of extinction due to the rapid loss of its habitat due to climate change and human activities (Zhang and Kong 2019) (Figure 1). *Betula microphylla* generally grows in wetlands within the arid and semi-arid areas where evaporation is much higher than precipitation. Many populations seem to have gone extinct during the past decades. For example, in the Caotanhu wetland, *B. microphylla* was historically abundant (Zhang and Kong 2019) and also existed in 2011, but the population had disappeared in 2018 (pers. comm.). A decrease in the level of groundwater due to climate change and human activities resulting in high salinity is thought to be responsible for the extinction of *B. microphylla* in the Caotanhu wetland (Li et al. 2019; Zhang and Kong 2019).

**Figure 1.**
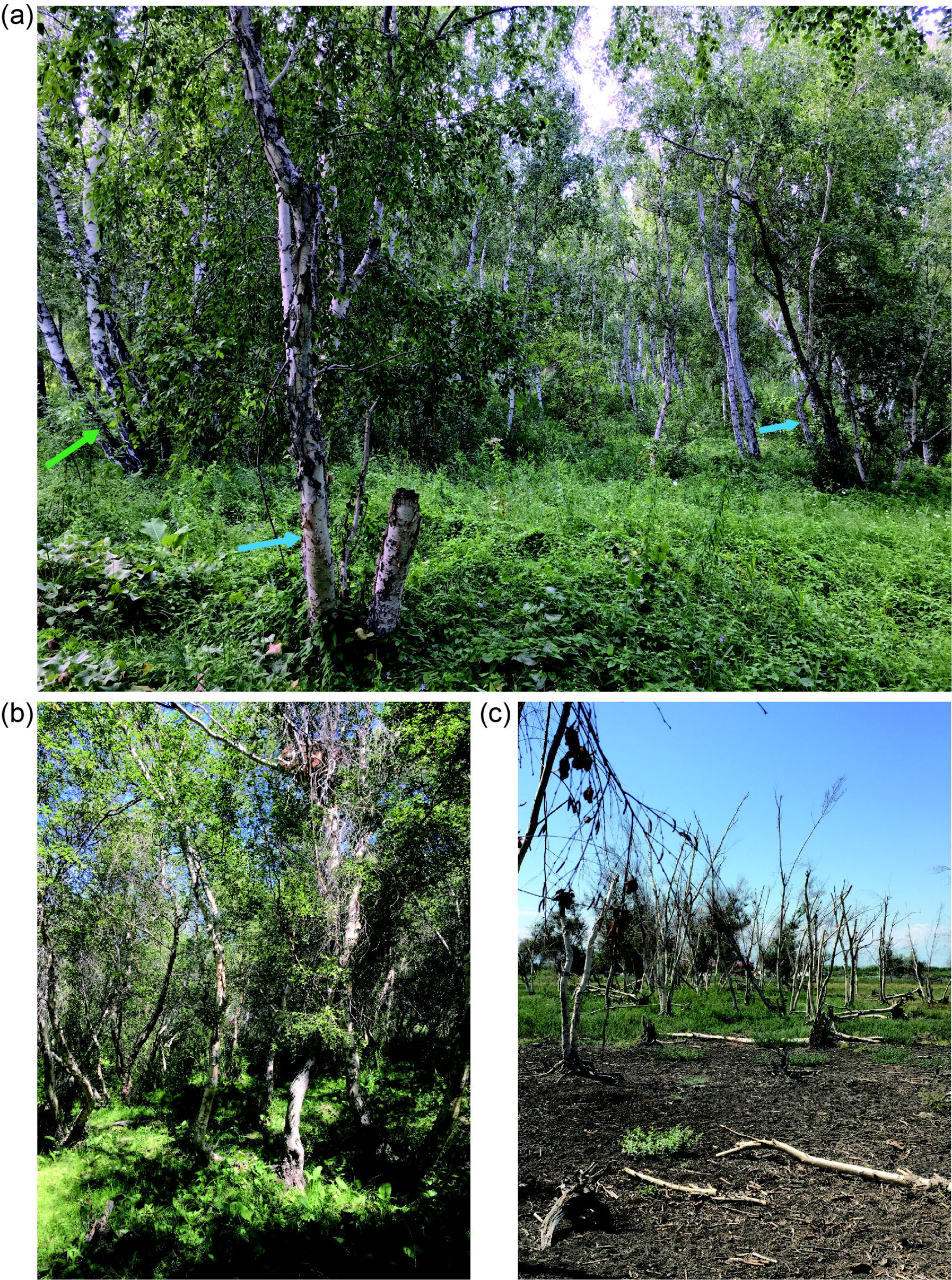
Habitat of *B. microphylla* at populations (a) YBW, (b) HBK and (c) MGH. Blue and orange arrows indicate *B. platyphylla* and *B. microphylla*, respectively.

*Betula microphylla* has an overlapping range with the more common *B. tianshanica* and both species are tetraploids (Ashburner and McAllister 2016). *Betula microphylla* is mainly distributed in the Altay Mountains (to the north of Xinjiang province). Historical occurrence data shows it used to occur in the Tianshan Mountains where *B. tianshanica* is common. The two tetraploid species form a morphological continuum with no clear boundaries (Figure S1), hindering conservation efforts. In Xinjiang province, several varieties of *B. microphylla* have been proposed based on minor morphological differences, such as *B. microphylla* var. *ebinurica*, which was named according to the locality where it was discovered (Yang et al. 2006). Several studies have suggested that these varieties need to be conserved urgently (Duan et al. 2018; Yang et al. 2006). However, no systematic classification is available to guide species conservation. The ranges of *B. microphylla* and *B. tianshanica* also overlap with that of the widespread species *B. platyphylla* but as the latter is diploid, any hybrids with the tetraploid species are unlikely to be viable.

Here, we used SSRs and restriction-site associated DNA sequencing (RAD-seq) to investigate the population genetic structure of *B. tianshanica* and *B. microphylla* and assess evidence for inter-specific genetic admixture. Combining marker sets with different ascertainment biases can enable us to better distinguish between alternative evolutionary scenarios, such as ancient or recent hybridisation and incomplete lineage sorting (Borrell et al. 2018). We also characterised local genetic diversity in *B. platyphylla* to check for any hybridisation with our two focal species. Our specific questions are: (1) Do *B. microphylla, B. tianshanica* and *B. platyphylla* form distinct genetic clusters? (2) Does hybridisation occur among *B. microphylla, B. tianshanica* and *B. platyphylla?* (3) If hybridisation occurs, does it potentially pose a threat to *B. microphylla?* To this end, we genotyped 265 individuals of *B. microphylla* and *B. tianshanica* collected from ten populations at 15 microsatellite markers and we conducted RAD-seq for 49 individuals with four or five individuals randomly chosen from each population. We genotyped 42 *B. platyphylla* individuals obtained from three populations using the microsatellite markers.

## 2. MATERIALS AND METHODS

### 2.1 Field sampling

Populations were located based on past records and samples were collected in July and August of 2018. Within each population, samples were separated by at least 20 meters to avoid sampling of the same clone. Sampling locations were recorded using a GPS system (UniStrong). A herbarium specimen was created for each individual and these were dried in a plant press. A total of five populations were from the Tianshan Mountains, and five were from the Altay Mountains. Populations SLM, EYQ and EYQE from the Tianshan Mountains are treated as *B. tianshanica* and populations HBK, TC, JMN, YBW and XY from the Altay Mountains treated as *B. microphylla*. Population ABH is tentatively defined as *B. microphylla* var. *ebinurica* (Yang et al. 2006) and population MGH tentatively defined as *B. microphylla* var. *microphylla* (pers. comm.), respectively. In addition, 42 samples of *B. platyphylla* were collected with 23, 17 and two individuals collected from population XEXL, YBW and XY, respectively. In total, we collected 136 samples from the Tianshan Mountains and 129 samples from the Altay Mountains and 42 *B. platyphylla* samples from the Altay Mountains.

### 2.2 Ecological niche modelling

To determine the potential distribution of *B. tianshanica, B. microphylla* and *B. platyphylla*, we sourced coordinate points from our fieldwork and literatures. A total of 23 coordinate points for *B. tianshanica*, 10 for *B. microphylla* and 26 for *B. platyphylla* from Xinjiang were obtained with points being at least 5 km apart from each other. Nineteen bioclimatic layers were downloaded from the WorldClim database (http://www.worldclim.org/) at a 2.5 arc-min resolution, for the LGM (CCSM v4 scenario), MID (CNRM-CM5 scenario), current (the period 1960-1990) and future periods (2050 and 2070, using MIROC-ESM scenario). Highly correlated variables (correlation coefficient >0.7) were identified using ‘raster.cor.matrix’ function within the R package ‘ENMTools’ version 0.2 (https://github.com/danlwarren/ENMTools) and were removed to avoid overfitting, with six retained for analysis. These six variables with low correlation were BIO1 (annual mean temperature), BIO2 (mean diurnal range), BIO4 (temperature seasonality), BIO9 (mean temperature of driest quarter), BIO12 (annual precipitation) and BIO15 (precipitation seasonality). Ecological niche model (ENM) was developed for each species in MaxEnt (Phillips et al. 2006). We performed 50 randomly subsampled replicate runs with 25% of observations retained for cross-validation. Models were further evaluated using the receiver operating characteristic (ROC) and area under curve (AUC). The AUC value greater than 0.9 indicates a very good prediction, while greater than 0.7 means the predicted model is better than a random model (Swets 1988). To compute niche overlap among the three species, we calculated Schoener’s D and Hellinger distance I (Schoener 1970; Warren et al. 2008) using the ‘raster.overlap’ function of the R package ‘ENMTools’ version 0.2.

### 2.3 Morphometric analysis

Five intact and mature leaves from each pressed specimen were selected for morphometric analysis. In total, we scanned 1325 leaves from the 265 samples using a Hewlett-Packard printer (LaserJet Pro MFP M128fn) with a resolution of 600 d.p.i. We selected thirteen landmarks from each scanned leaf as described in our previous study (Hu et al. 2019). Briefly, the coordinates of each of the thirteen landmarks were recorded on digitized leaves and were converted to a configuration of 26 cartesian coordinates in 13 pairs (x, y) for each leaf using the program ImageJ (Abràmoff et al. 2004). Principal component analyses (PCAs) on the normalized matrix were conducted on the level of each leaf and each specimen, respectively, using the program MORPHOJ (Klingenberg 2011).

### 2.4 DNA extraction and SSR genotyping

Genomic DNA was isolated from cambium tissues following a previously modified 2× CTAB (cetyltrimethylammoniumbromide) protocol (Wang et al. 2013). The quality of isolated genomic DNA was assessed with 1.0 % agarose gels and then was diluted to a concentration of 10-20 ng/ul for microsatellite genotyping. Fifteen microsatellite loci developed for *B. pendula* (Kulju et al. 2004), *B. pubescens* ssp. *tortuosa* (Truong et al. 2005), *B. maximowicziana* (Tsuda et al. 2009) and *B. platyphylla* var. *japonica* (Wu et al. 2002) were used for genotyping our samples (Table S1). The 5’ terminus of the forward primers was labeled with FAM, HEX or TAM fluorescent probes. Each microsatellite locus was amplified individually prior to being artificially combined into four multiplexes (Table S1). In order to avoid errors caused by size overlapping, loci with significant length differences were labeled using the same dye. PCR procedures were provided in supplementary data.

Microsatellite alleles were scored using the software GENEMARKER 2.4.0 (Softgenetics) and checked manually. Individuals with more than three missing loci were excluded, resulting in 307 individuals in the final dataset.

### 2.5 SNP genotyping

A total of 50 DNA samples were delivered to KEGENE Company for library preparation using restriction enzymes *PstI* and *MspI* to digest DNA. Libraries were sequenced with an Illumina HiSeq 2500 using pair-end 150-bp sequencing. Adapter sequences were removed using a custom Perl script. The raw data were cleaned using the process_radtags module of Stacks v2.2 (Catchen et al. 2013) in paired-end mode. A sliding-window step was performed with a window size of 15% of the length of the read. Reads with a required quality of below 30 within the sliding-window and unpaired reads were discarded. Clean reads of the sequenced samples were aligned to the whole genome of *B. pendula* (Salojärvi et al. 2017) using BWA-MEM v.0.7.17-r1188 algorithm in BWA (v0.7.17) with default parameters (Li and Durbin 2009). Reads with non-specific matches were discarded. One individual was *B. platyphylla* and excluded for subsequent analyses, resulting in 49 samples.

Alignments were converted from sequence alignment map (SAM) format to sorted, indexed binary alignment map (BAM) files (SAMtools v1.8) (Li et al. 2009). The MarkDuplicates tool from the Genome Analysis Tool Kit (GATK) (v 4.1.4.) was used to mark duplicates (DePristo et al. 2011; McKenna et al. 2010). The HaplotypeCaller from GATK was used to call genotypes for each sample (v4.1.4), generating intermediate gVCF files. The CombineGVCFs tool was used to merge these intermediate gVCF files into a combined VCF file, which used as the input file for joint genotyping using the GenotypeGVCFs tool. SNPs were filtered using a mapping quality (MP) threshold of 40, a variant confidence (QUAL) of 30, a normalized QUAL score (QD) of 2, a maximum symmetric odds ratio (SOR) of 3, a minimum depth (DP) of 5 and a maximum depth of 200, a maximum probability of strand bias (FS) and excess heterozygosity (ExcessHet) of 60 and 54.69, respectively, and a minimum Z-score of read mapping qualities (MQRankSum) and position bias (ReadPosRankSum) of −12.5 and −8.0, respectively. The SelectVariants filtering tool was applied to select SNPs present in at least 50% of the samples.

### 2.6 Population structure analysis of SSRs and SNPs

Principal coordinate (PCO) analysis was performed on SSR data of *B. microphylla, B. tianshanica* and *B. platyphylla* using POLYSAT (Clark and Jasieniuk 2011) implemented in R 3.6.1 (R Core Team 2019), based on pairwise genetic distances calculated according to Bruvo et al. (Bruvo et al. 2004). For nucleotide SNPs a principal component analysis (PCA) was carried out using the ‘adegenet’ R package 2.1.1 (Jombart 2008), setting the ploidy level as four.

Both datasets were analyzed in STRUCTURE 2.3.4 (Pritchard et al. 2000) to identify the most likely number of genetic clusters (K) with a ploidy of four. Ten replicates of the STRUCTURE analysis were performed for SSRs and five replicates for SNPs due to computational limitations. We used 1,000,000 iterations and a burn-in of 100,000 for each run at each value of K from 2 to 10 for SSRs and 100,000 generations and a burn-in of 100,000 for each run (five replicates) for SNPs. For both datasets we used the admixture model, with an assumption of correlated allele frequencies among populations. Individuals were assigned to clusters based on the highest membership coefficient averaged over the ten independent runs. The number of genetic clusters that make most biological sense was estimated using both “Evanno test” (Evanno et al. 2005) using the program Structure Harvester (Earl and vonHoldt 2012) and the “Thermo-dynamic Integration” (TI) method (Verity and Nichols 2016) for microsatellite data. As for SNPs, we did not use the TI method as it was not designed for thousands of SNPs. Replicate runs were grouped based on a symmetrical similarity coefficient of >0.9 using the Greedy algorithm in CLUMPP (Jakobsson and Rosenberg 2007) and visualized in DISTRUCT 1.1 (Rosenberg 2004). Gene diversity (Nei 1987) and allelic richness (El Mousadik and Petit 1996) were calculated in the software FSTAT 2.9.4 (Goudet 1995) using the microsatellite dataset. The tetraploid genotypes were treated as two diploid individuals as described by Tsuda et al. (2017) and Hu et al. (2019). Isolation by distance (IBD) analysis was performed for *B. tianshanica* and *B. microphylla*.

### 2.7 Comparison of genetic admixture

We ploted the estimated level of genetic admixture derived from STRUCTURE analyses of SSRs against RAD-seq data at the optimal K value. Then we conducted Mann–Whitney U-test to test the difference between genetic admixture estimated using the different marker sets using the function ‘wilcox. test’. All statistical analyses were conducted in R 3.6.1 (R Core Team 2019).

## 3. RESULTS

### 3.1 Predicted distributions

The constructed ecological niche models performed well for *B. tianshanica* with AUC ranged from 0.932 to 0.940 and SD from 0.025 to 0.029 for the five time-scales and were satisfactory for *B. microphylla* with AUC ranged from 0.813 to 0.839 and SD from 0.044 to 0.065. The most important environmental predictors were annual precipitation for *B. tianshanica* and precipitation seasonality for *B. microphylla*. In the present day, suitable habitat for *B. tianshanica* at the LGM is predicted to be limited and may exist in the western part of the Tianshan Mountains which is outside Xinjiang whereas suitable habitat for *B. microphylla* is widespread in north Xinjiang and on the west of Xinjiang (Figure 2). During the Holocene, suitable habitat for *B. tianshanica* was widespread along the Tianshan Mountians and for *B. microphylla* was widespread in the Altay Mountains and its environs. Suitable habitat for *B. tianshanica* seems to have been stable from the Holocene to the present and is predicted to be much larger by 2050 and 2070, with a shift northwards. However, suitable habitat for *B. microphylla* has contracted since the Holocene and is predicted to become very limited and fragmented by 2050 and 2070 (Figure 2). Analysis of range overlap at a conservative occurrence probability threshold (0.40) showed very little overlap between *B. tianshanica* and *B. microphylla* (I=0.036, Schoener’s D=0.001) in the LGM and more overlap during the Holocene (I=0.229, Schoener’s D=0.115) and at the present (I=0.257, Schoener’s D=0.125). Range overlap is predicted to be larger between *B. tianshanica* and *B. microphylla* in 2050 (I=0.362, Schoener’s D=0.303) and 2070 (I=0.349, Schoener’s D=0.298) (Figure 2). Suitable habitat for *B. platyphylla* seems to have been stable from the Holocene to the present and is predicted to become very limited to the Altay Mountains by 2050 and 2070. Range overlap between *B. microphylla* and *B. platyphylla* is consistently broader than between *B. tianshanica* and *B. platyphylla* during all these periods (Figure 2).

**Figure 2.**
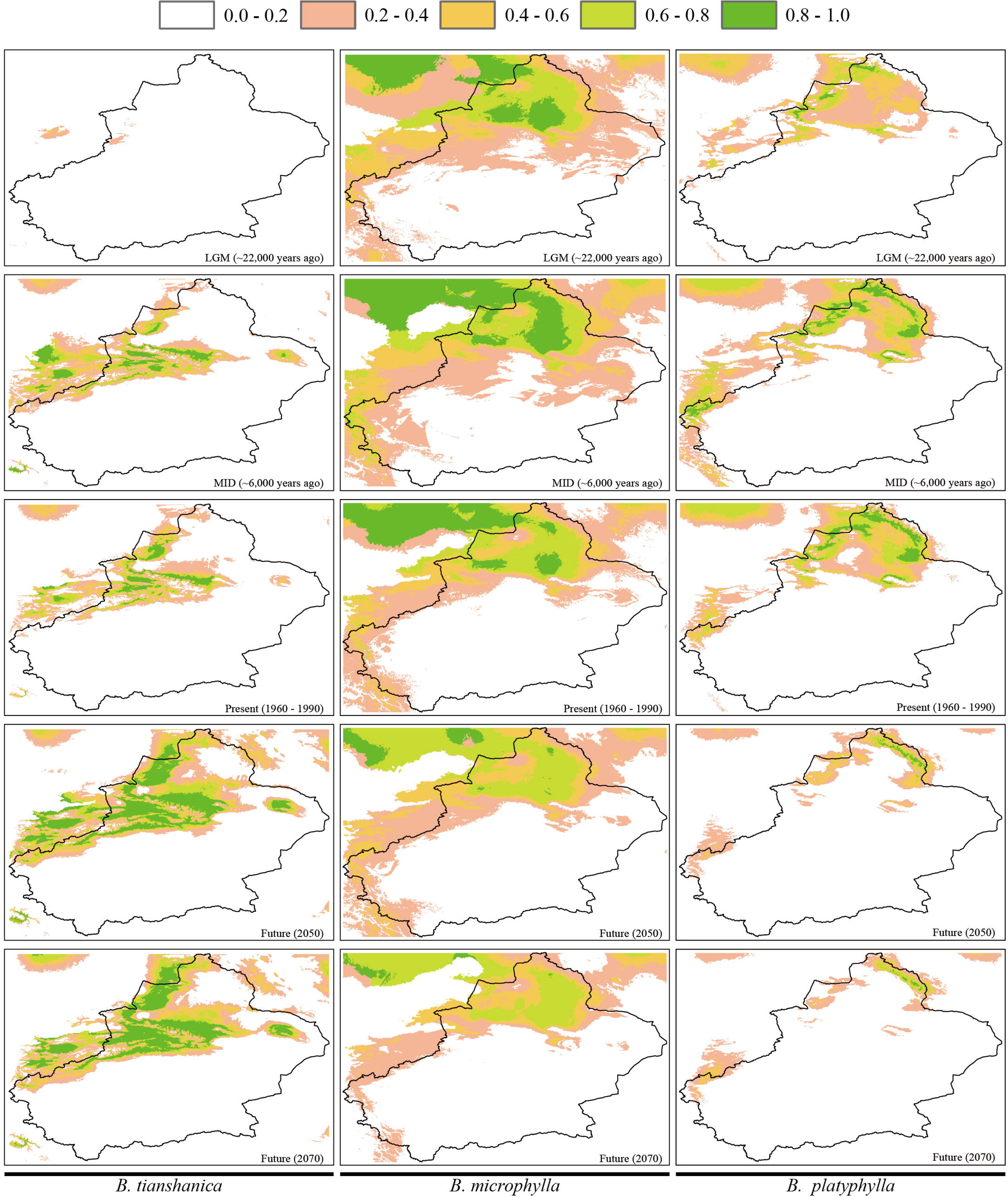
The predicted suitable habitat for *B. tianshanica, B. microphylla* and *B. platyphylla* during LGM, the Holocene, at present and in the future (2050, 2070).

### 3.2 Morphometric analysis

PCA on leaf morphology of all collected samples of *B. microphylla* and *B. tianshanica* revealed a single overlapping cluster. PC1 and PC2 explained 30.3% and 18.7% of the variation among leaves, respectively (Figure 3) and summarized 40.9% and 23.4% of the variation among trees, respectively (Figure S2).

**Figure 3.**
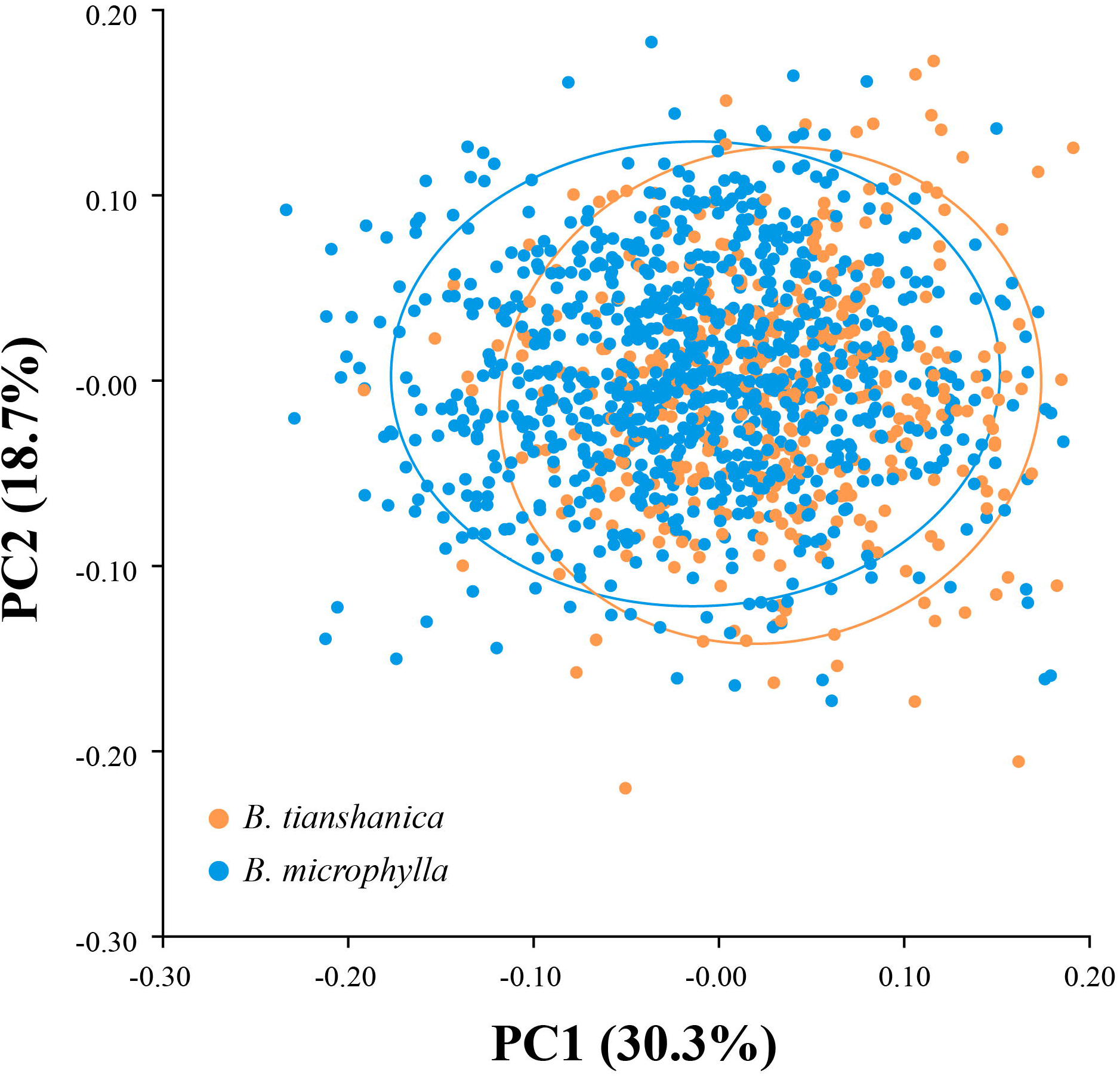
PCA analysis performed among leaves. Each orange and blue dot represents each leaf of *B. tianshanica* and *B. microphylla*, respectively.

### 3.3 SSR analyses

PCO analysis based on Bruvo’s genetic distances among all these samples revealed three clusters (Figure 4) in coordinates 1 and 2 that between them accounted for 26.7% of the total variation. *Betula platyphylla* was well differentiated from the other species, whereas the tetraploid species formed separate clusters that were closer together, with some individuals bridging the gap between them.

**Figure 4.**
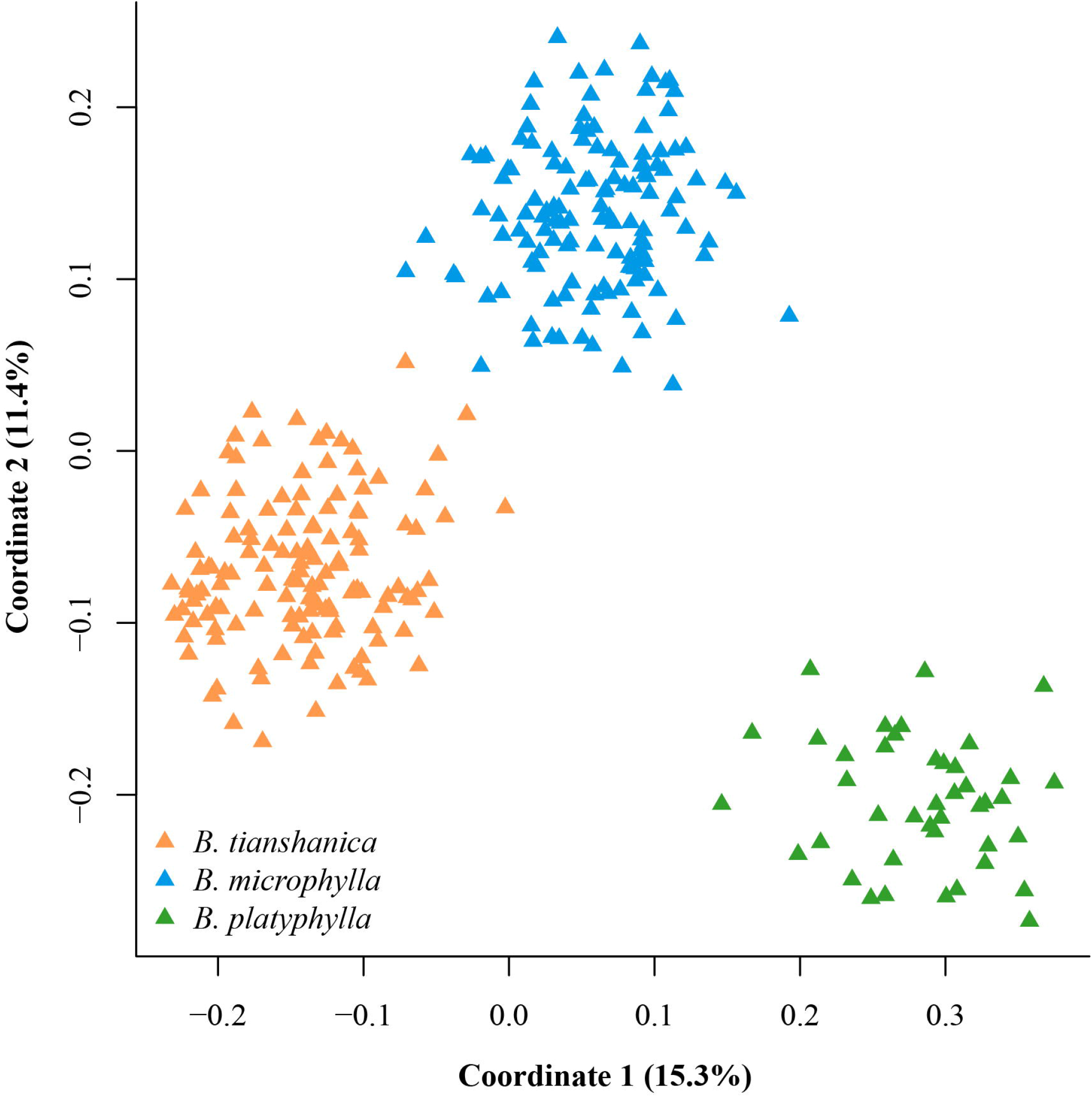
Principal component analysis of *Betula* samples at 15 SSRs.

Consistent with these PCO results, our STRUCTURE analyses identified three clusters. The “TI” method showed K = 3 was the optimal value whereas ΔK criterion failed to identify the optimal K value (Figure S3ab). Cluster I, which is shaded orange in Figure 5, included populations EYQE, EYQ, SLM, ABH and MGH, which correspond with the Tianshan Mountains and *B. tianshanica*. Cluster II, which is shaded blue in Figure 5, included populations HBK, TC, JMN, YBW and XY which correspond to *B. microphylla* from the Altay Mountains. Cluster III, which is shaded green in Figure 5, corresponded with *B. platyphylla* (Figure 5a). *Betula tianshanica* populations had a relatively low level of genetic admixture from *B. microphylla*, but there seemed to be one early-generation hybrid between *B. tianshanica* and *B. microphylla* in population SLM. *Betula microphylla* populations TC and HBK had a relatively high level of genetic admixture from *B. tianshanica* whereas populations YBW and XY had a low level of genetic admixture. Limited genetic admixture was detected from *B. platyphylla* into sympatric populations of *B. microphylla* but little admixture was found from *B. platyphylla* into *B. tianshanica* (Figure 5a).

**Figure 5.**
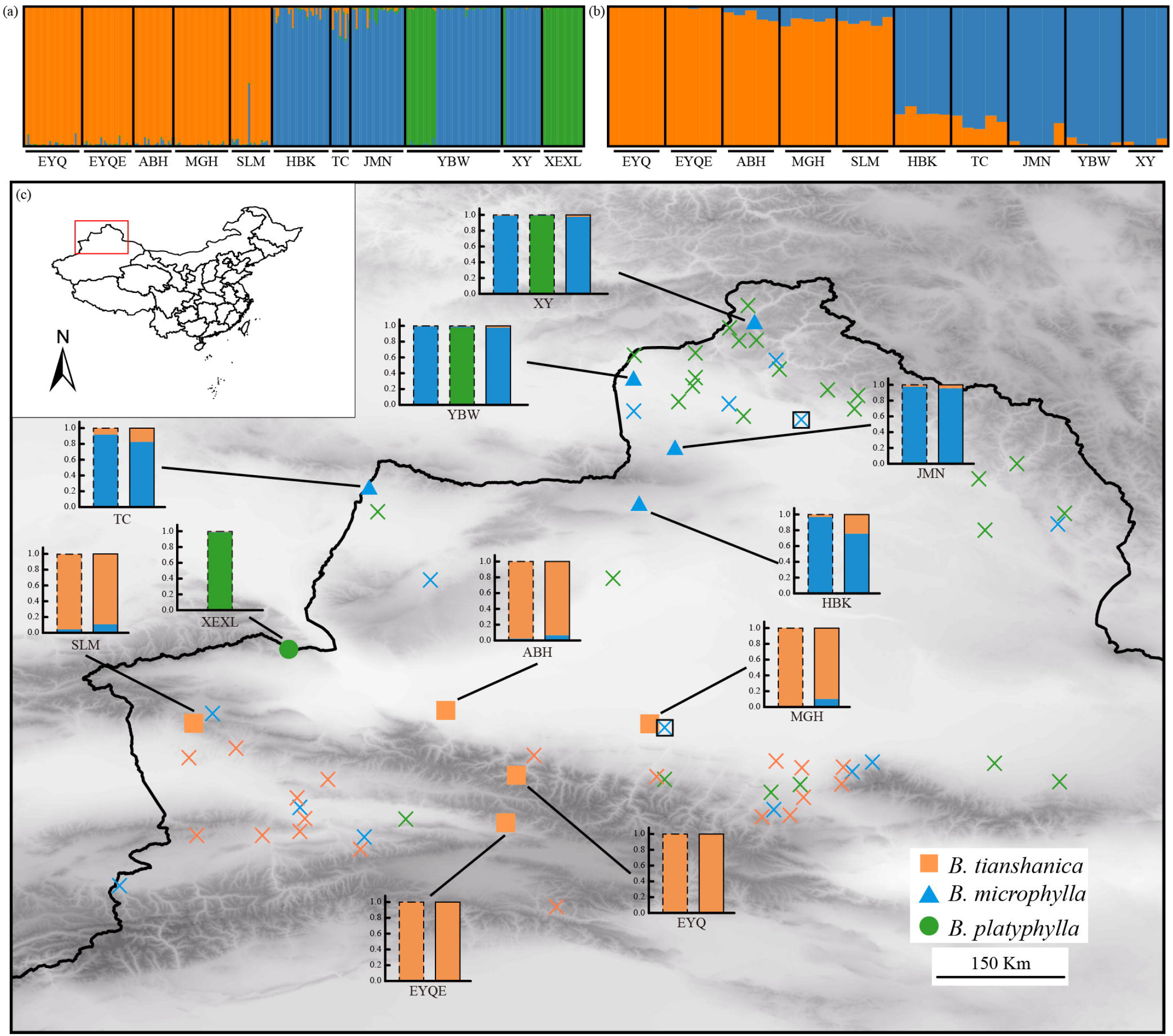
STRUCTURE results at K=3 and K=2 based on SSRs (a), and SNPs (b), respectively and a map showing sampling sites (c). Orange circle, blue triangle and green dot represent *B. tianshanica, B. microphylla*, and *B. platyphylla* respectively. ABH and MGH indicate populations where species were regarded as *B. microphylla* var. *ebinurica* and *B. microphylla* var. *microphylla*, respectively, but were *B. tianshanica*. Orange, blue and green crosses indicate records of *B. tianshanica, B. microphylla* and *B. platyphylla*, respectively.

Populations ABH and MGH have previously been regarded as containing the sub-species *B. microphylla* var. *ebinurica* and *B. microphylla* var. *microphylla*. Our result suggests that these sub-species are mainly *B. tianshanica* at the genetic level, with introgression from *B. microphylla* (Figure 5a).

We found no significant signal of IBD within populations of *B. tianshanica* and *B. microphylla* but significant signal of IBD between *B. tianshanica* and *B. microphylla* (Figure S4). Gene diversity and allelic richness were marginally higher for *B. microphylla* than *B. tianshanica* (Table 1).

**Table 1.**
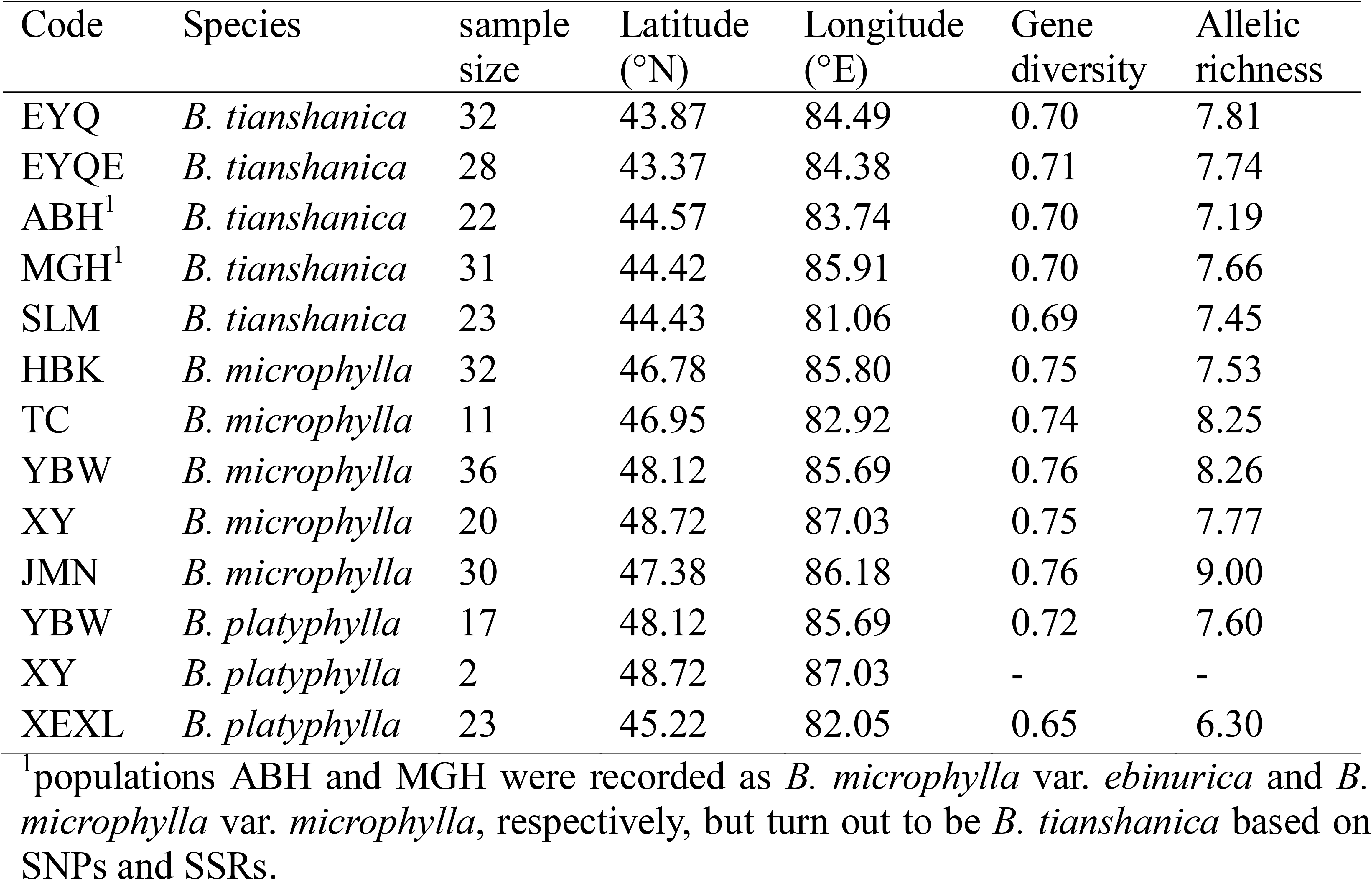
Sampling information and genetic diversity of ten populations of *B. tianshanica, B. microphylla* and *B. platyphylla* based on SSRs.

| Code | Species | sample size | Latitude (°N) | Longitude (°E) | Gene diversity | Allelic richness |
| --- | --- | --- | --- | --- | --- | --- |
| EYQ | *B. tianshanica* | 32 | 43.87 | 84.49 | 0.70 | 7.81 |
| EYQE | *B. tianshanica* | 28 | 43.37 | 84.38 | 0.71 | 7.74 |
| ABH<sup>1</sup> | *B. tianshanica* | 22 | 44.57 | 83.74 | 0.70 | 7.19 |
| MGH<sup>1</sup> | *B. tianshanica* | 31 | 44.42 | 85.91 | 0.70 | 7.66 |
| SLM | *B. tianshanica* | 23 | 44.43 | 81.06 | 0.69 | 7.45 |
| HBK | *B. microphylla* | 32 | 46.78 | 85.80 | 0.75 | 7.53 |
| TC | *B. microphylla* | 11 | 46.95 | 82.92 | 0.74 | 8.25 |
| YBW | *B. microphylla* | 36 | 48.12 | 85.69 | 0.76 | 8.26 |
| XY | *B. microphylla* | 20 | 48.72 | 87.03 | 0.75 | 7.77 |
| JMN | *B. microphylla* | 30 | 47.38 | 86.18 | 0.76 | 9.00 |
| YBW | *B. platyphylla* | 17 | 48.12 | 85.69 | 0.72 | 7.60 |
| XY | *B. platyphylla* | 2 | 48.72 | 87.03 | - | - |
| XEXL | *B. platyphylla* | 23 | 45.22 | 82.05 | 0.65 | 6.30 |

<sup>1</sup>populations ABH and MGH were recorded as *B. microphylla* var. *ebinurica* and *B. microphylla* var. *microphylla*, respectively, but turn out to be *B. tianshanica* based on SNPs and SSRs.

### 3.4 SNP analyses

The number of raw reads per sample ranged from 6,436,408 to 92,924,972 and the number of trimmed reads per sample ranged from 5,827,435 to 84,297,044 with between 5,165,913 and 78,364,760 mapped to the *B. pendula* reference genome for each sample. The individual read mappings resulted in 64.1% to 93.7% of mapped reads per individual. Over 90% of trimmed reads mapped successfully for 42 out of 49 samples are (Table S2). After filtering (> one million raw reads, only SNVs, coverage between 5 and 200, <50% missing data), 386,363 variant sites were found present in at least one individual and 982 were present in all 49 individuals. As the basis of our population analyses we used variants present in at least 50% of individuals, of which 35,814 were biallelic, 1364 were triallelic and 39 were tetra-allelic.

PCA analyses (Figure S5) based on genotype calls for these 37,217 loci in 49 individuals indicated two clusters corresponding to *B. microphylla* and *B. tianshanica*. STRUCTURE analysis (Figure 5b) also supported an optimal K of 2, according to the ΔK criterion (Figure S6). In *B. tianshanica*, little introgression from *B. microphylla* was detected in populations EYQ and EYQE, with the highest admixture levels of 0.4% and 1.5%, respectively (Figure S7). More introgression from *B. microphylla* was estimated in populations SLM, ABH and MGH, with highest admixture values of 12.5%, 9.5% and 13.8%, respectively (Figure 5b, Figure S7). In *B. microphylla* more introgression was found from *B. tianshanica*. Admixture values of 28.1% and 22.9% were estimated for HBK and TC, respectively and a lower level of admixture from *B. tianshanica* was detected in the more northerly populations with highest admixture values of 16.6%, 6.4% and 5.6% in populations JMN, YBW and XY, respectively (Figure 5b, Figure S7).

### 3.5 Comparison of genetic markers

Both STRUCTURE analyses of SNPs and SSRs distinguished *B. microphylla* from *B. tianshanica* (Figure S8). SNPs indicated higher introgression from *B. tianshanica* to *B. microphylla* than did SSRs (P=0.0296, Mann–Whitney U test). Whereas for introgression from *B. microphylla* to *B. tianshanica* both SNPs and SSRs (P=0.1386, Mann–Whitney U test) yielded similar estimates (Figure S8). The estimated admixture from *B. tianshanica* to *B. microphylla* was higher for populations HBK and TC based on SNPs than based on SSRs (Figure 6a). Similarly, the estimated admixture from *B. microphylla* to *B. tianshanica* was higher for populations MGH and SLM based on SNPs than based on SSRs (Figure 6b).

**Figure 6.**
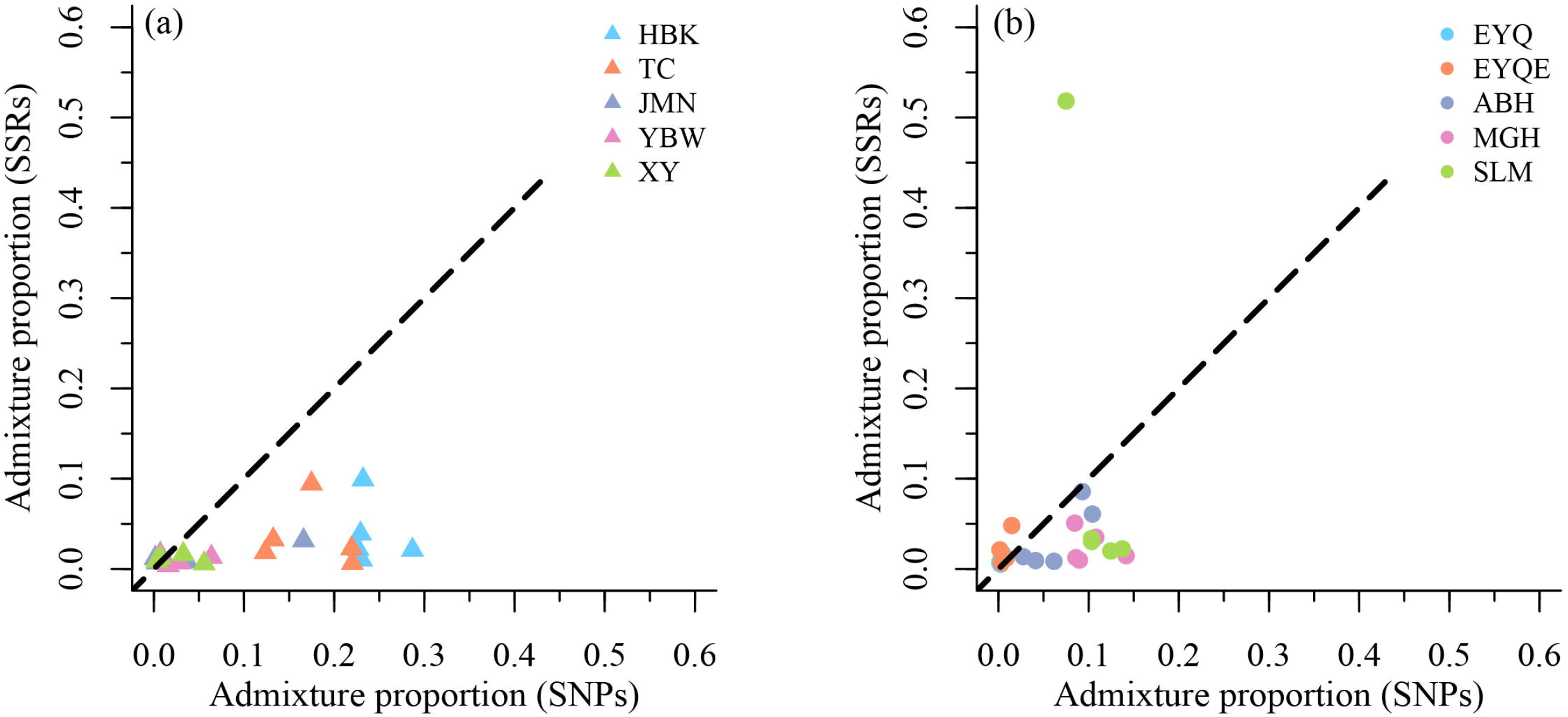
Comparison of the genetic admixture using SNPs and SSRs. (a) and (b) indicate genetic admixture from *B. microphylla* into *B. tianshanica* and genetic admixture from *B. tianshanica* into *B. microphylla*, respectively.

## 4. DISCUSSION

### 4.1 Species delimitation

Our genetic analyses identified two distinct clusters corresponding to the rare species *B. microphylla* and the more common species *B. tianshanica*, providing strong evidence that they are genetically distinct entities despite the continuum of leaf morphology between them (Ashburner and McAllister 2016). We found very rare populations, previously referred to as *B. microphylla* var. *ebinurica* and *B. microphylla* var. *microphylla*, to be mainly *B. tianshanica* at the genetic level with genetic admixture from *B. microphylla*. A hybrid history of these populations seems plausible as geographically they occur between the main current ranges of *B. tianshanica* and *B. microphylla*, in the watershed of the Tianshan Mountains. Previous research noted that *B. microphylla* var. *ebinurica* resembled both *B. microphylla* and *B. tianshanica* (Yang et al. 2006). On the basis of our genetic findings, we suggest that *B. microphylla* var. *ebinurica* and *B. microphylla* var. *microphylla* from MGH should not be treated as sub-species, but as *B. tianshanica*.

### 4.2 Unusual patterns of two-way introgression

Both SNP and SSR markers identified bi-directional introgression between *B. tianshanica* and *B. microphylla*, with greater apparent introgression from *B. tianshanica* into *B. microhylla* than vice versa. The SNP markers tended to show higher levels of introgression than SSRs (Figure 5ab), a pattern also found in a comparison of SNPs and SSRs by Bradbury et al. (Bradbury et al. 2015) but not in other similar studies (Bohling et al. 2019; Zohren et al. 2016). One individual, namely, SLM020, was estimated to have 51.8% admixture from *B. microphylla* based on SSRs but have 7.5% admixture from *B. microphylla* based on SNPs (Figure 5ab; Figure S8). This may be explained by the smaller sample size of SSRs, which may be less representative of the whole genomes.

Introgression into *B. tianshanica* (SLM and MGH) tended to be higher in areas where occurrence of *B. microphylla* has been reported in the past. Introgression into *B. microphylla* tended to be higher in proximity to *B. tianshanica* populations, but also appeared to have occurred in areas without records of *B. tianshanica* populations. The apparent presence of *B. tianshanica-derived* alleles to the north of its current range could be explained by hybridisation involving unexpectedly high pollen flow or undiscovered populations of *B. tianshanica* in the region. It could also be due to selection for adaptive introgression, and populations of *B. microphylla* could be adapting to a changing climate using alleles from *B. tianshanica*. Another explanation for the more widespread *B. tianshanica* alleles could be incomplete lineage sorting (ILS) but the geographic signals of genetic admixture (Barton 2001) that we found suggest that ILS is at most making a minor contribution to the patterns of apparent admixture.

### 4.3 Threats to *B. microphylla*

Our study suggests that declining populations of *B. microphylla* face threats from hybridisation with the more common *B. tianshanica*. As our ENM results predict a northward expansion of *B. tianshanica* in the future, we expect hybridisation to become even more common between the two species than it is currently. In our field observations, *B. tianshanica* was usually found growing along riversides or on mountain slopes adjacent to rivers so it may be that its northward advance is helped by seed dispersal via rivers flowing north from the Tianshan Mountains.

The threat of hybridisation is in addition to the already known threat of habitat degradation and loss to *B. microhylla*. During our fieldwork we observed no seedlings of *B. microphylla* in the populations where we collected samples, which may provide further evidence for habitat degradation. Without recruitment *B. microphylla* populations cannot survive. Populations of *B. microphylla* in the northern regions generally grow in wetlands, surrounded by vastly inhospitable arid or semi-arid lands, thus limiting the successful dispersal of pollens or seeds. Overgrazing and crop plantation are modifying these habitats greatly, posing unprecedented risk to the existence of both *B. microphylla* and *B. tianshanica*. For example, the MGH population which is only 20 km away from Shihezi city is facing extinction due to human activities.

### 4.4 Implications for conservation and management

Accurate species delimitation is crucial for conservation. Here, both SSRs and SNPs distinguished *B. tianshanica* from *B. microphylla*, providing a basis for conservation of the latter. Unexpectedly, *B. microphylla* var. *ebinurica* (ABH population) and *B. microphylla* var. *microphylla* (MGH population) are for the first time identified as *B. tianshanica* with some introgression from *B. microphylla*. Given this, we think it is unnecessary to prioritise these populations for conservation (cf. Yang et al., 2006; Duan et al., 2018; Li et al., 2019). The genetic integrity of *B. microphylla*, appears to be threatened by gene flow from *B. tianshanica*, and could be contributing further to the decline of *B. microphylla*. More research is needed to understand if this gene flow is beneficial or detrimental to *B. microphylla* in the context of climate change. If it is beneficial it could be enhanced by assisted gene flow, but if it is detrimental, action may be needed to maintain genetically pure *B. microphylla* populations. Meanwhile, it would be wise to conserve and expand northerly populations of *B. microphylla* with little admixture from *B. tianshanica*.

## ACKNOWLEDGEMENTS

We thank Dr. Laura Kelly, from Royal Botanic Gardens Kew for valuable comments on the manuscript and Dr. Jihong Huang from Research Institute of Forest Ecology, Environment and Protection, Chinese Academy of Forestry for sharing coordinate points of *B. microphylla* with us. This work was funded by the National Natural Science Foundation of China (31770230 and 31600295) and Funds of Shandong ‘Double Tops’ Program (SYL2017XTTD13).

## AUTHOR CONTRIBUTION

NW and DH conceived the project. DH, NW and LW collected samples. JD, LW, ZL and FW carried out lab work. JD analyzed the results. NW wrote the initial draft, RJAB and JSB provided substantial input thereafter.

---

**Figure S1** Pairs of leaves from a subset of *B. tianshanica* and *B. microphylla* samples in the present study, showing upper and lower sides. The first eight samples are *B. tianshanica* and the remaining samples are *B. microphylla*.

**Figure S2** PCA analysis performed among samples. Blue and red dots represent samples of *B. tianshanica* and *B. microphylla*, respectively.

**Figure S3** The best number of clusters inferred using the “Thermo-dynamic Integration” method (a) and “Evanno test” method (b).

**Figure S4** Isolation by distance (IBD) analyses for *B. tianshanica, B. microphylla* and between *B. tianshanica* and *B. microphylla*.

**Figure S5** PCO analysis based on SNPs.

**Figure S6** The output of Structure Harvester showing that K = 2 is the optimal value based on SNPs.

**Figure S7** Admixture value for each population of *B. tianshanica* and *B. microphylla* based on SNPs.

**Figure S8** An alignment of STRUCTURE results at K = 2 for SNPs and SSRs, respectively.

## Table legends

**Table S1.** Details of microsatellite primers used in the present study.

**Table S2.** Detailed information about results of RAD-seq for each sample in this study.

## Notes

### Competing Interest Statement

The authors have declared no competing interest.

