## Supplemental Data for "Bidirectional introgression between *Betula tianshanica* and *Betula microphylla* and its implications for conservation"

### *New Phytologist* Supporting Information

The following Supporting Information is available for this article:

**Figure S1** Pairs of leaves from *B. tianshanica* and *B. microphylla*.

**Figure S2** PCA analysis performed among samples.

**Figure S3** The best number of clusters.

**Figure S4** Isolation by distance (IBD) analyses.

**Figure S5** PCO analysis based on SNPs.

**Figure S6** The output of Structure Harvester.

**Figure S7** Admixture value for each population.

**Figure S8** An alignment of STRUCTURE results.

**Table S1** Details of microsatellite primers used in the present study.

**Table S2** Detailed information about results of RAD-seq.

**Methods S1** Procedures of PCR amplification.

**Figure S1** Pairs of leaves from a subset of *B. tianshanica* and *B. microphylla* samples in the present study, showing upper and lower sides. The first eight samples are *B. tianshanica* and the remaining samples are *B. microphylla*.

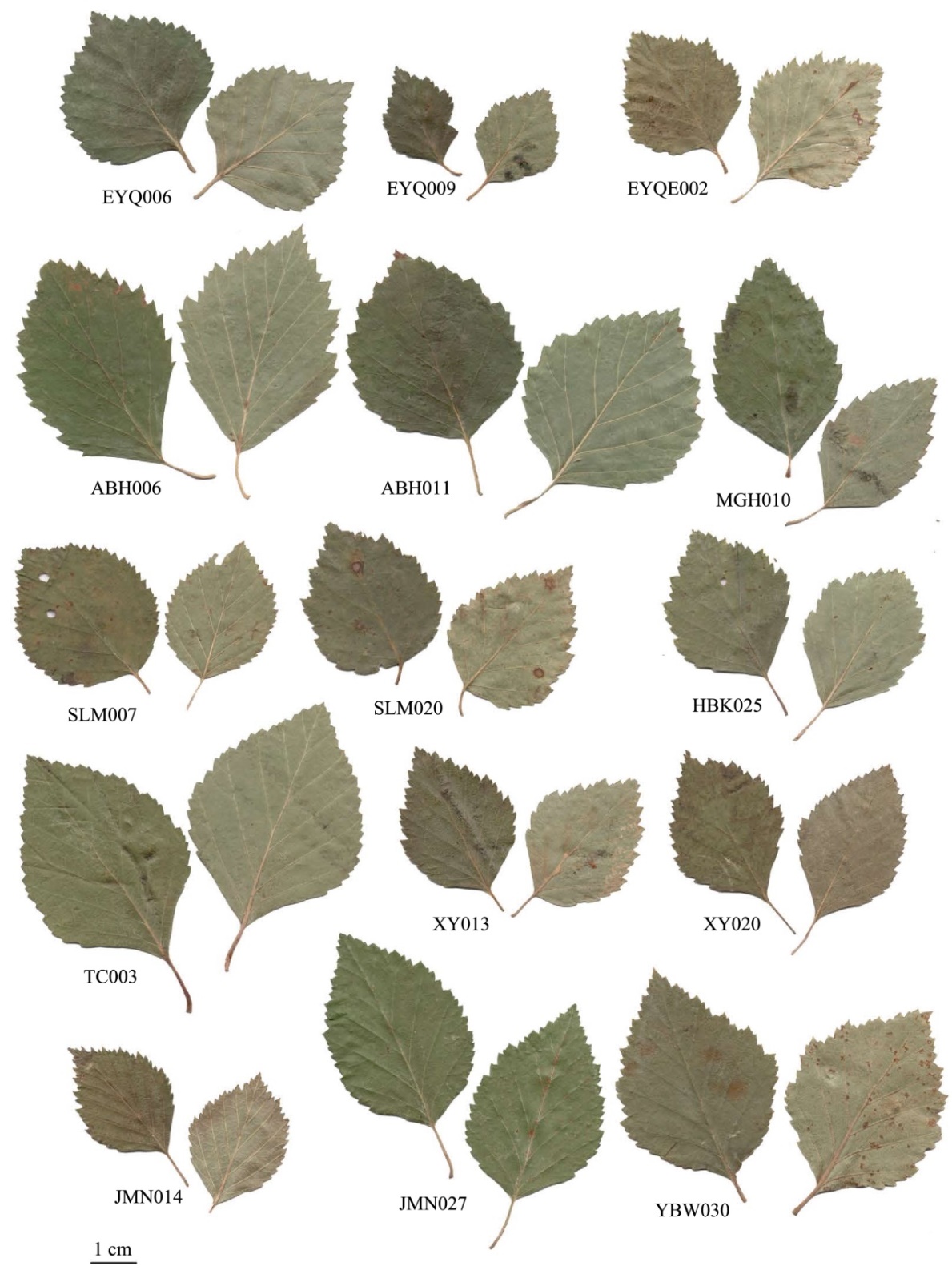

**Figure S2** PCA analysis performed among samples. Blue and red dots represent samples of *B. tianshanica* and *B. microphylla*, respectively.

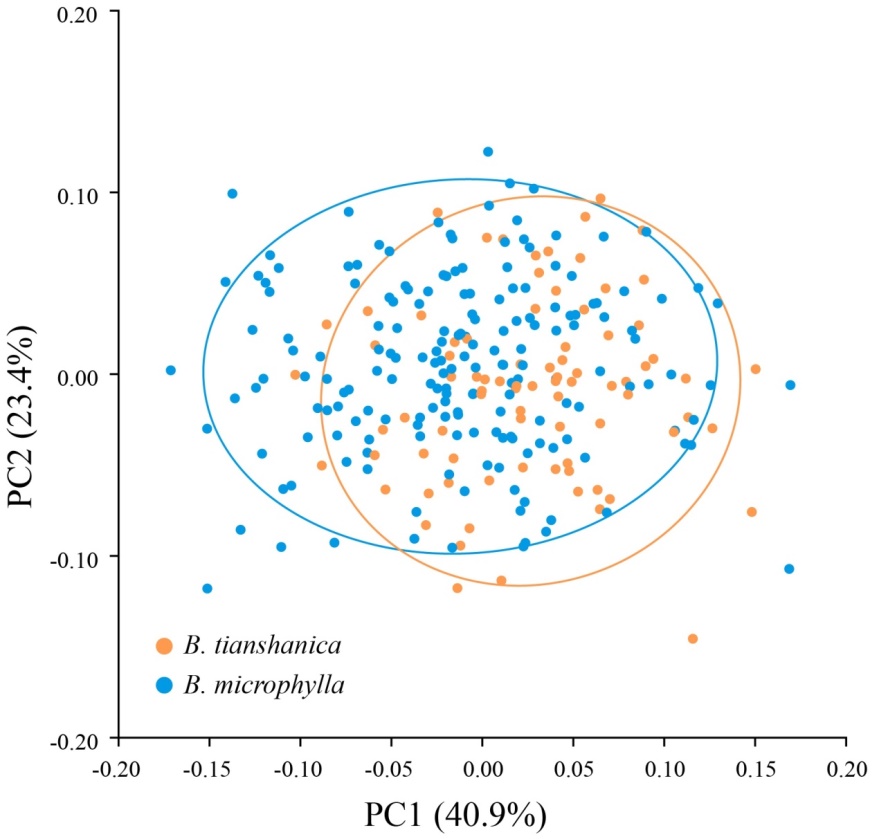

**Figure S3** The best number of clusters inferred using the “Thermo-dynamic Integration” method (A) and“Evanno test” method (B).

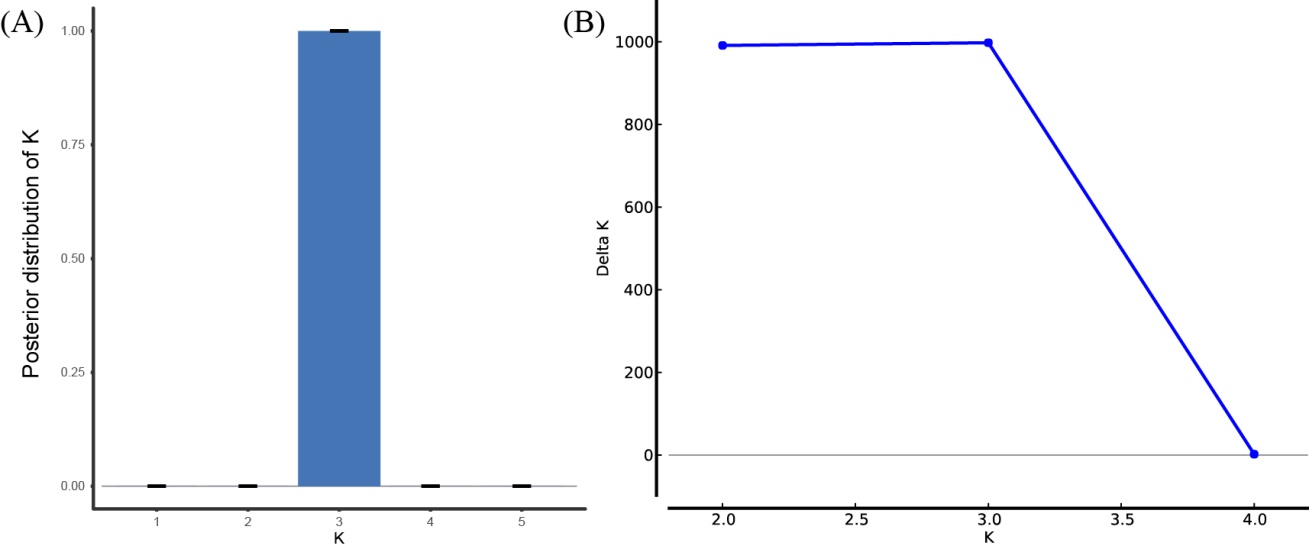

**Figure S4** Isolation by distance (IBD) analyses for *B. tianshanica*, *B. microphylla* and between *B. tianshanica* and *B. microphylla*.

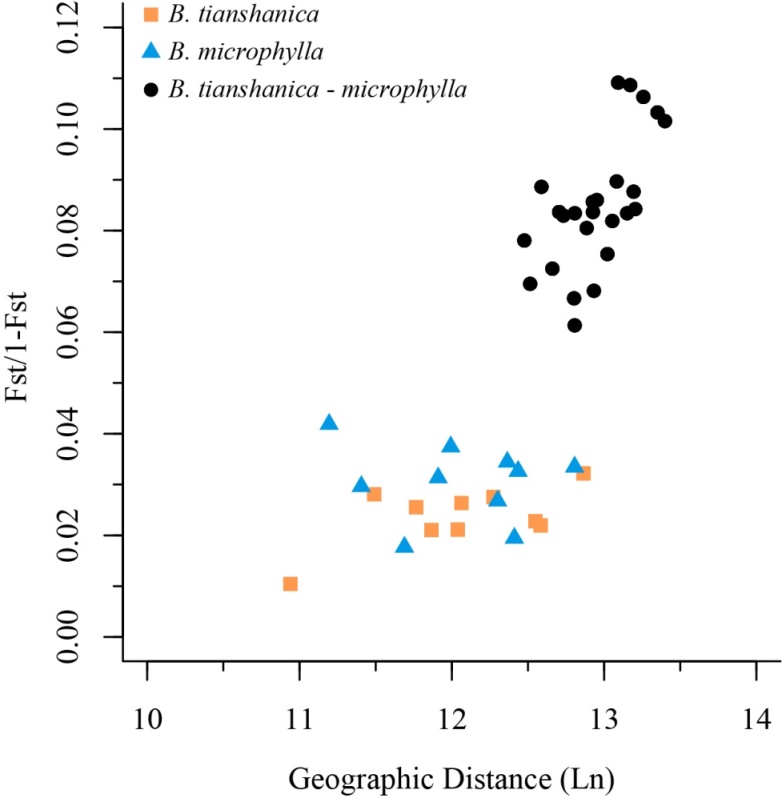

**Figure S5** PCO analysis based on SNPs.

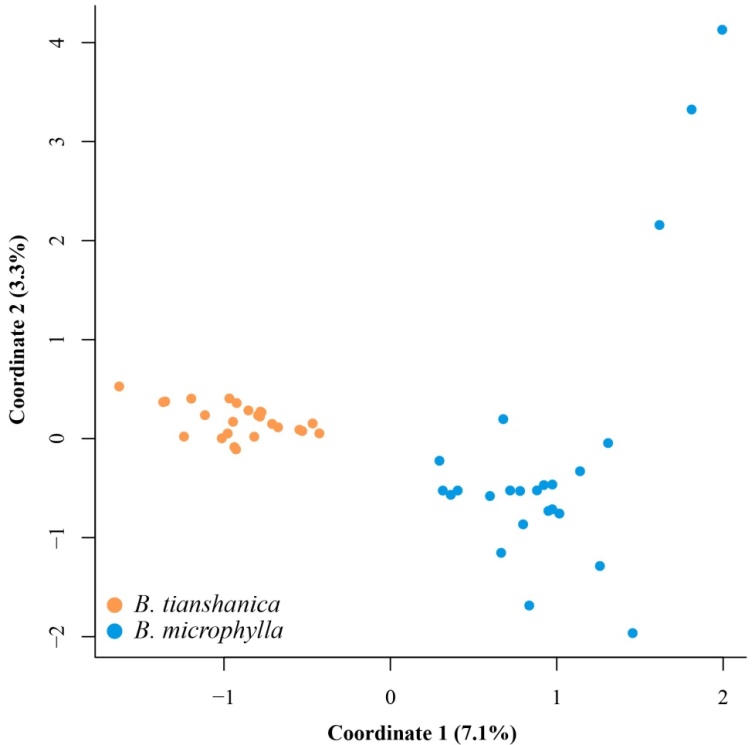

**Figure S6** The output of Structure Harvester showing that K = 2 is the optimal value based on SNPs.

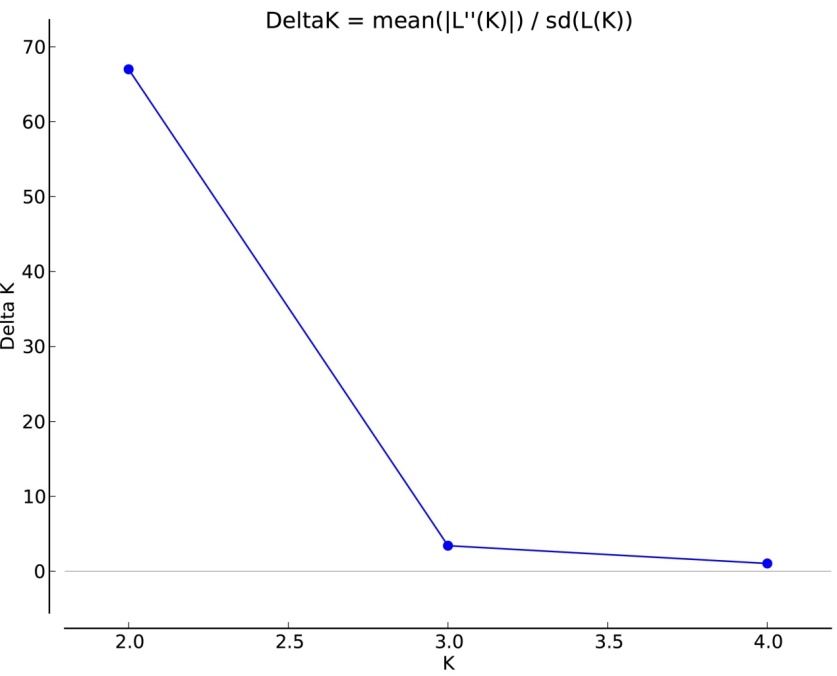

**Figure S7** Admixture value for each population of *B. tianshanica* and *B. microphylla* based on SNPs.

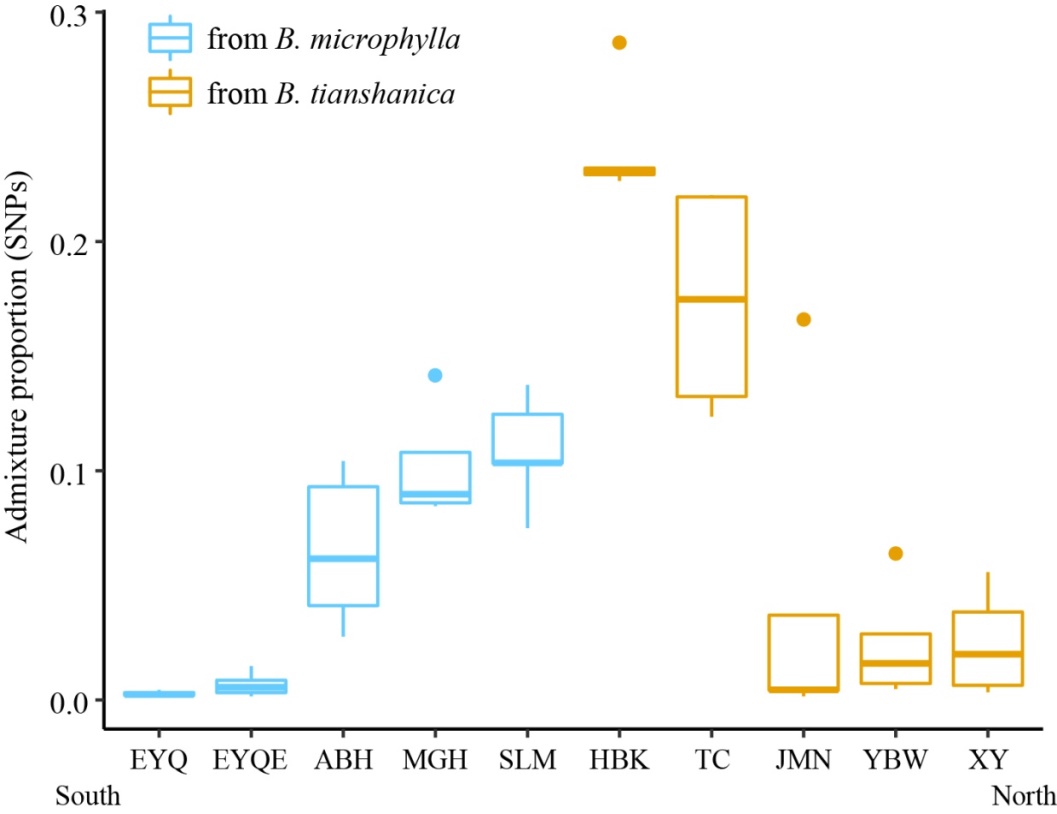

**Figure S8** An alignment of STRUCTURE results at K = 2 for SNPs and SSRs, respectively.

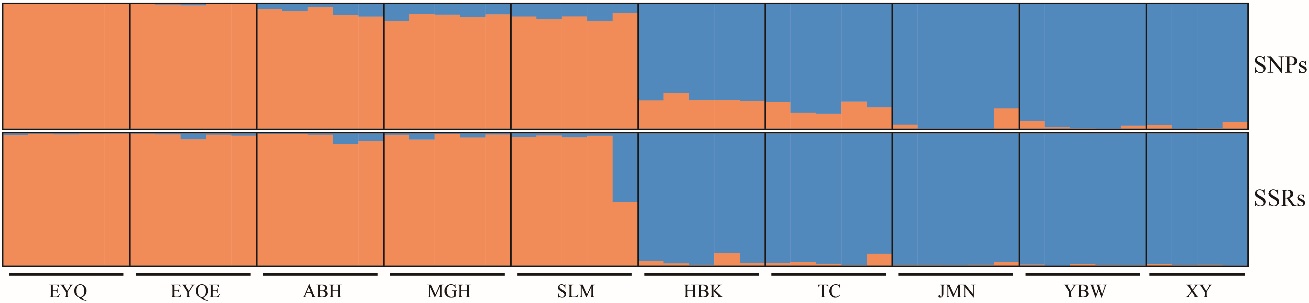

**Table S1** Details of microsatellite primers used in the present study.

| Locus | Dye | Primer sequence (5´-3´) | Allele Size (bp) | Repeat | Multiplex |
| --- | --- | --- | --- | --- | --- |
| CD277117 | TAMRA | F:AGGGGACCCACTAATTTTTCTGT R:GTTTCTGCTTCAAGGCTTCCCAT | 111–130 | (AC)_4_TTTTTCCGGGTGCGTTA(CT)_3_G(TC)_6_ | Multiplex 1 |
| L7.1 | FAM | F: GTTTTGGGTTTCCACTTCCA  R: ACTGGTAATACCTTTACCAAGCC | 146-152 | (CT)_12_CCTT(CT)_4_ |  |
| L1.10 | HEX | F: ACGCTTTCTTGATGTCAGCC  R: TCACCAAGTTCCTGGTGGAT | 168–209 | (GA)_4_AA(GA)_10_ |  |
| CD277113 | HEX | F: ACAATATCTCACAAATCTGCCGC  R:GTTTAACCTGAACGTCCTCAAAGGTCC | 272–278 | (TC)_9_ |  |
| L2.7 | HEX | F: CCGCCGGTAACACTAAACC  R: GAGGGAAGAAAATTCAACGG | 141–186 | (TC)_8_(TA)_8_(TG)_11_TT(TG)_3_ | Multiplex 2 |
| L3.1 | TET | F:CTCCTTAGCTGGCACGGAC  R:CCCTTCTTCATAAAACCCTCAA | 219–241 | (CT)_3_CC(CT)_2_CC(CT)_13_AT(CT)_5_ |  |
| L2.3 | TAMRA | F: CAGTGTTTGGACGGTGAGAA  R: CGGGTGAAGTAGACGGAACT | 198–220 | (AG)_16_ |  |
| CD277839 | FAM | F:ATAACCGACGAGCTCAAGAAAGA  R:GTTTGCACCTCGAAACTGGTACAACA | 309–321 | (GAA)_6_ |  |
| Bp10 | HEX | F:GTTGTAATGCAAACACATGGG  R:TCTGTGTCATAATTGGGTAGG | 124-152 | (GT)_28_ | Multiplex 3 |
| Bo.F394 | TET | F:AATGCAGCATCTCTTACC  R:CACGCAATAATATGGAAA | 128–194 | (TC)_13_ |  |
| L7.3 | TAMRA | F:GGGGATCCAGTAAGCGGTAT  R:CACACGAGAGATAGAGTAACGGAA | 178–226 | (GT)_18_(GA)_14_ |  |
| L5.4 | TAMRA | F:AAGGGCACCTGCAGATTAGA  R:AAAATTGCAACAAAACGTGC | 230–262 | (TC)_26_ |  |
| Bp01 | TAMRA | F:GAAAGGATCTGTATAGCCAAC  R:ACCACATGGCAGCAATTCTAG | 130-164 | (GT)_10_ | Multiplex 4 |
| L022 | HEX | F:AACGGACAAATTCACGGGTA  R:GGAGTTCATGGATTGGAGGA | 172-214 | (CT)_18_ |  |
| L2.5 | HEX | F: CTATATTGGCTCCAAGCAC  R: ACACCCACACTGACAGATAA | 94–128 | (CT)_9_ |  |

**Table S2** Detailed information about results of RAD-seq for each sample in this study.

| Samples | Total reads | Clean reads | Mapped reads | Percentage (%) | Variants |
| --- | --- | --- | --- | --- | --- |
| ABH002 | 22,475,060 | 20,470,618 | 18,593,631 | 90.83 | 767,189 |
| ABH006 | 48,199,030 | 44,217,234 | 40,942,719 | 92.59 | 904,613 |
| ABH011 | 31,205,020 | 28,536,064 | 26,581,461 | 93.15 | 801,662 |
| ABH016 | 28,735,424 | 25,732,310 | 23,649,154 | 91.90 | 801,310 |
| ABH021 | 25,217,090 | 23,139,957 | 21,626,913 | 93.46 | 808,414 |
| EYQ006 | 33,827,760 | 30,947,088 | 28,865,643 | 93.27 | 842,950 |
| EYQ007 | 36,321,080 | 33,016,152 | 30,678,467 | 92.92 | 872,426 |
| EYQ008 | 63,654,644 | 57,693,888 | 53,536,370 | 92.79 | 919,625 |
| EYQ009 | 38,041,540 | 34,568,326 | 32,163,642 | 93.04 | 1,064,365 |
| EYQ010 | 40,113,388 | 36,757,992 | 34,456,542 | 93.74 | 915,437 |
| EYQE002 | 28,226,766 | 25,801,291 | 23,764,923 | 92.11 | 845,007 |
| EYQE007 | 35,812,068 | 32,185,086 | 29,236,920 | 90.84 | 696,493 |
| EYQE015 | 50,041,252 | 45,391,906 | 41,653,904 | 91.77 | 796,223 |
| EYQE023 | 55,191,428 | 49,817,099 | 42,699,595 | 85.71 | 1,158,599 |
| EYQE028 | 40,832,196 | 36,797,509 | 32,000,134 | 86.96 | 984,181 |
| HBK003 | 28,519,988 | 25,670,634 | 23,564,169 | 91.79 | 762,194 |
| HBK008 | 29,380,596 | 26,327,067 | 23,886,203 | 90.73 | 654,523 |
| HBK010 | 28,816,444 | 26,431,837 | 24,466,846 | 92.57 | 721,584 |
| HBK025 | 29,869,808 | 26,752,853 | 24,409,438 | 91.24 | 562,925 |
| HBK027 | 22,186,354 | 20,229,008 | 18,583,990 | 91.87 | 533,568 |
| JMN001 | 35,579,862 | 32,695,546 | 30,452,281 | 93.14 | 971,750 |
| JMN008 | 21,853,894 | 19,680,601 | 17,696,654 | 89.92 | 558,683 |
| JMN014 | 26,630,834 | 24,437,999 | 22,754,582 | 93.11 | 795,887 |
| JMN020 | 27,935,568 | 25,100,206 | 23,078,712 | 91.95 | 504,024 |
| JMN027 | 27,324,702 | 24,945,621 | 23,022,704 | 92.29 | 936,148 |
| MGH002 | 30,732,804 | 28,195,418 | 26,389,695 | 93.60 | 908,010 |
| MGH010 | 36,626,596 | 32,932,692 | 30,467,967 | 92.52 | 1,037,523 |
| MGH021 | 44,548,708 | 40,096,278 | 37,181,612 | 92.73 | 1,106,413 |
| MGH028 | 43,840,020 | 39,568,398 | 36,683,146 | 92.71 | 1,314,442 |
| MGH037 | 92,924,972 | 84,297,044 | 78,364,760 | 92.96 | 1,055,716 |
| SLM003 | 31,690,364 | 28,622,823 | 26,266,991 | 91.77 | 713,392 |
| SLM007 | 21,354,224 | 19,220,209 | 17,678,076 | 91.98 | 710,509 |
| SLM012 | 44,444,352 | 39,770,590 | 25,495,927 | 64.11 | 846,613 |
| SLM016 | 13,963,346 | 12,863,722 | 11,885,864 | 92.40 | 580,601 |
| SLM020 | 39,633,616 | 35,388,520 | 32,585,867 | 92.08 | 740,338 |
| TC001 | 30,503,082 | 27,845,577 | 25,103,999 | 90.15 | 896,341 |
| TC003 | 32,248,250 | 29,547,110 | 27,252,865 | 92.24 | 990,882 |
| TC005 | 56,769,696 | 51,511,103 | 47,552,745 | 92.32 | 1,415,106 |
| TC007 | 27,177,788 | 24,817,526 | 19,497,940 | 78.57 | 659,457 |
| TC009 | 37,170,352 | 33,765,308 | 31,104,110 | 92.12 | 841,425 |
| XY002 | 18,962,698 | 17,330,321 | 15,843,680 | 91.42 | 715,306 |
| XY005 | 6,436,408 | 5,827,435 | 5,165,913 | 88.65 | 386,363 |
| XY013 | 26,756,152 | 24,280,670 | 22,224,887 | 91.53 | 813,667 |
| XY020 | 24,598,622 | 22,583,411 | 19,574,453 | 86.68 | 933,633 |
| YBW008 | 49,876,272 | 44,911,464 | 41,097,222 | 91.51 | 1,167,660 |
| YBW022 | 39,899,880 | 35,867,916 | 33,104,182 | 92.29 | 1,012,843 |
| YBW030 | 31,592,088 | 28,821,818 | 26,847,910 | 93.15 | 771,562 |
| YBW043 | 35,906,472 | 32,667,622 | 30,251,138 | 92.60 | 795,400 |
| YBW052 | 48,663,100 | 44,203,230 | 40,958,112 | 92.66 | 804,656 |

**Methods S1** Procedures of PCR amplification.

Polymerase chain reactions (PCRs) were conducted for each of the microsatellites using Tiangen master mix (Tiangen Biotech, Beijing, China). The final reaction volume was 10 μL, including 5 μL Tiangen PCR Master Mix, 0.15-0.2 μL of primers (10 μm each in initial volume), 3.6 μL H_2_O and 5–20 ng of DNA dissolved in 1.0 μL TE buffer. An initial denaturation step at 94 °C for 30 s was followed by 30 cycles of denaturation (94 °C for 30 s), annealing (57 °C for 90 s) and extension (72 °C for 60 s) steps, and a final extension step at 60 °C for 30 min. Fragment lengths were determined by capillary gel electrophoresis using an ABI 3730xl capillary.
